# Precision Genome and Base Editing by Regulating mRNA Nuclear Export Using Selective Inhibitors of Nuclear Export (SINEs)

**DOI:** 10.1101/2021.04.17.440169

**Authors:** Yan-ru Cui, Shao-jie Wang, Tiancheng Ma, Peihong Yu, Jun Chen, Taijie Guo, Genyi Meng, Biao Jiang, Jiajia Dong, Jia Liu

## Abstract

CRISPR-based genome engineering tools are associated with off-target effects that constitutively active Cas9 protein may instigate. In the present study, we screened for irreversible small molecule off-switches of CRISPR-Cas9 and discovered that selective inhibitors of nuclear export (SINEs) could inhibit the cellular activity of CRISPR-Cas9 by interfering with the nuclear export of Cas9 mRNA. We subsequently found that SINEs, including an FDA-approved anticancer drug KPT330, could improve the specificities of CRISPR-Cas9-based genome and base editing tools in human cells.

## Main

Clustered Regularly-Interspaced Short Palindromic Repeats (CRISPR) and CRISPR-associated (Cas) proteins (CRISPR-Cas) are the bacterial immune system for defending bacteriophage infection^1^. Type II CRISPR contains a single streamlined nuclease that can be reprogrammed for various genome engineering applications^2–4^. In human cells, CRISPR-Cas9 can induce DNA double-strand breaks (DSBs) that are repaired by two competing mechanisms: non-homologous end joining (NHEJ) and, in the presence of DNA templates, homology-directed repair (HDR)^5^. Therapeutic genome editing often involves constitutively expressed Cas9 protein^6^, which may introduce excessive DSBs and error-prone NHEJ, leading to off-target mutations^7–9^, chromosomal rearrangement^10^ or genotoxicity^11^.

Recently, CRISPR-derived base editors^12^ (BEs) have been developed to overcome the adverse effects associated with CRISPR-based genome editing tools^13^. BEs are fusion proteins comprising a catalytically inactive Cas nuclease, a nucleobase deaminase and, in some cases, DNA glycosylase inhibitors^12,14^. BEs can generate nucleotide substitutions without introducing DSBs or DNA template^12^ and are thus optimal choice for precision gene therapy^15,16^. Similar to genome editing tools, uncontrolled BEs are also found to be associated with high frequency of off-target events^17–19^. This problem is particularly recognized in CBEs in comparison with relatively high-fidelity ABEs^20,21^.

To mitigate these side effects, temporal control-enabling CRISPR-Cas inhibitors have been investigated. Thus far, naturally occurring, phage-derived anti-CRISPR proteins (Acrs) are most characterized CRISPR inhibitors. A number of Acrs have been identified for type I^22–26^, type II^27–29^ and type V^26,30^ CRISPR-Cas systems. These inhibitors exert their functions by disrupting distinct steps of CRISPR-Cas actions such as single-guide RNA (sgRNA) binding^31^, DNA binding^32^ or DNA cleavage^32^. In nature, Acrs are used by bacteriophages to escape the CRISPR-Cas immunity in bacteria^22,33^. In genome engineering applications, Acrs are adapted to modulate CRISPR-Cas functions in a variety of host cells including bacteria^34^, yeast^35^ and mammalian cells^27,32,34,36,37^. In addition, Acrs can be coupled with CRISPR-Cas system to construct biosensors^38^ or synthetic circuits in eukaryotes^39^. The ability of Acrs to achieve temporal and spatial^40^ or optogenetic^41^ control of CRISPR-Cas9 has enabled their applications for improving the targeting specificity^37^, cell type-dependent activity^42^ and cytotoxicity^11,43^ of CRISPR-based genome editing tools.

In addition to protein-based inhibitors, oligonucleotides^44^ and phage-derived peptides^31^ have also been developed as CRISPR off-switches. Importantly, small-molecule inhibitors of CRISPR-Cas9 have been identified using an *in vitro* high-throughput screening assay^45^. These small molecules can reversibly inhibit the cellular activity of CRISPR-Cas9 by disrupting Cas9-DNA interactions^45^. Despite of the myriad types and mechanisms, these CRISPR-Cas inhibitors remain poorly understood for their therapeutic potential, particularly the safety in human.

In the present study, we used an EGFP reporter-based live cell assay to screen for irreversible small-molecule inhibitors of CRISPR-Cas9. We discovered that selective inhibitors of nuclear export (SINEs), including a marketed anti-cancer drug KPT330 (selinexor), could regulate the cellular activity of CRISPR-Cas9 by interfering with the nuclear export of Cas9 mRNA. We subsequently found that these SINEs could be used to modulate the activity and specificity of CRISPR-Cas9-based genome editing, base editing and prime editing tools.

## Results

### Screening for irreversible small-molecule inhibitors of CRISPR-Cas9

While most existing CRISPR-Cas inhibitors exert their functions via reversible, non-covalent interactions, we speculated that irreversible inhibitors could be also suitable options for CRISPR-Cas9 that is exogenous to human cells. Therefore, we focused screening on a compact collection of approximately 500 small-molecule compounds with irreversible warheads. The screen was performed on a HEK293-based EGFP reporter cell line^46^ that carries an out-of-frame EGFP gene, the expression of which can be restored upon CRISPR-Cas9 targeting (Fig. 1a). Inhibition of CRISPR-Cas9 cellular activity by inhibitors will lead to reduced fraction of activated EGFP cells (Supplementary Fig. 1). The first round of screening identified several Michael acceptor-bearing compounds that exhibited efficient inhibition of EGFP activation. The second round of screening was performed on a focused library of marketed or investigational compounds containing Michael acceptors (Fig. 1b). Several candidate CRISPR-Cas9 inhibitors were identified with half maximal inhibitory concentrations (IC_50_) of less than 20 μM (Supplementary Table 1). Notably, an FDA-approved anticancer drug KPT330^47,48^ displayed greatest potency for Cas9 inactivation (Fig. 1b).

**Fig. 1:**
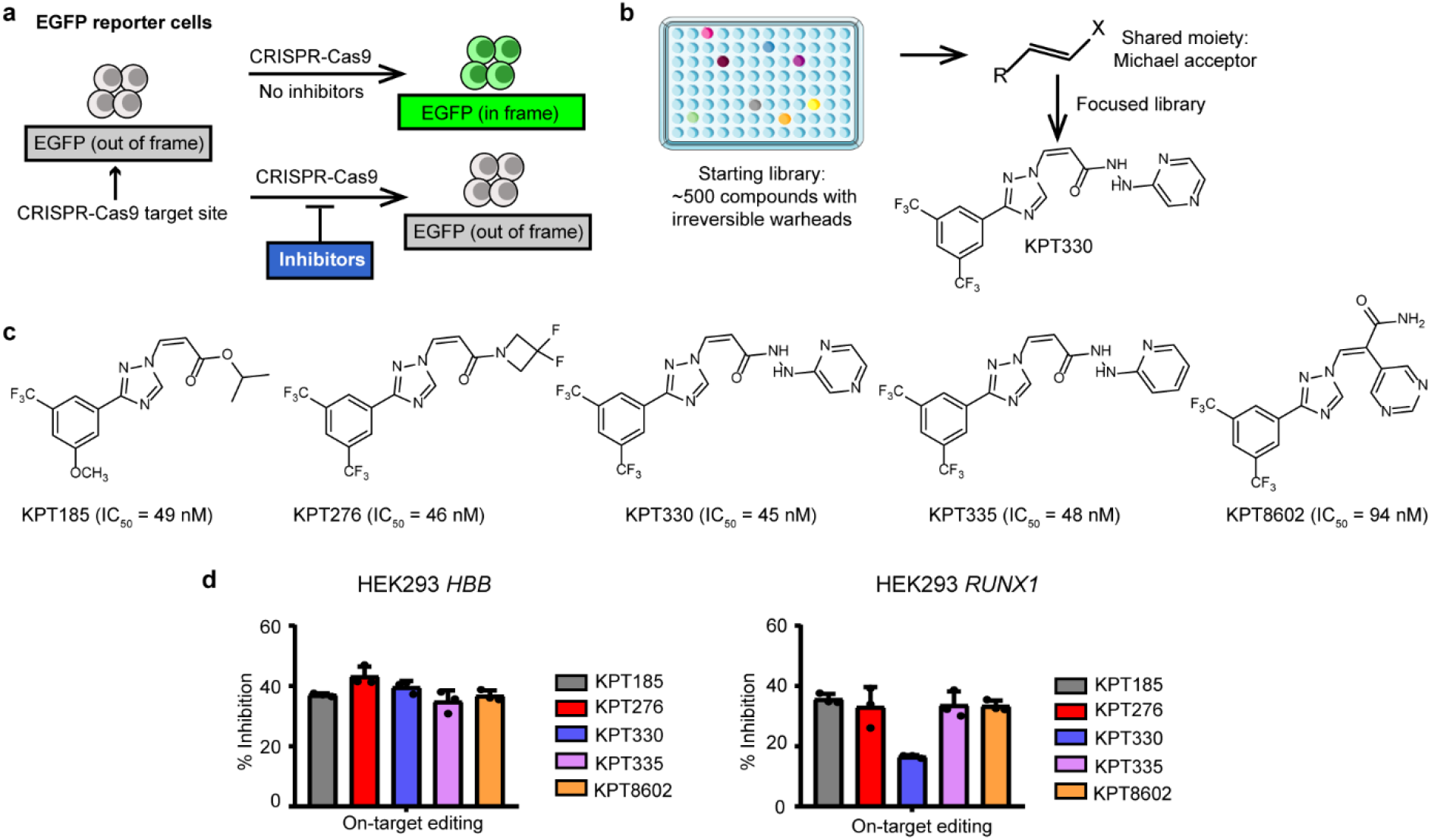
Identification of SINEs as CRISPR-Cas9 inhibitors. **a** Schematic presentation of EGFP reporter cell-based screening of CRISPR-Cas9 inhibitors. **b** Flowchart of the screening process. X, electron withdrawing group. **c** Potency of SINEs on inhibiting CRISPR-activated EGFP fluorescence. **d** SINEs at 0.5 μM inhibit the on-target editing activity of CRISPR-Cas9 at *HBB and RUNX1* genes in HEK293 cells. The data are shown as mean ± SD (*n* = 3).

Inspired by the results with KPT330, we investigated other selective inhibitors of nuclear export (SINEs) including KPT185, KPT276, KPT335 and KPT8602 for Cas9 inhibition. It was found that these SINEs could efficiently suppress CRISPR-Cas9-induced EGFP activation (Fig. 1c and Supplementary Fig. 2a). To examine the inhibitory effects of KPT330 on CRISPR-Cas9 at the endogenous genomic sites, we designed four sgRNAs targeting to human *EMX1*, *AAVS1* and *HBB* genes respectively. Similar to the procedure in EGFP activation assay, 10 μM KPT330 was added into the medium following the transfection of sgRNA- and Cas9-encoding plasmids in HEK293T cells. T7E1 assay analysis confirmed the efficient inhibition of CRISPR-Cas9 by KPT330 at endogenous genomic sites (Supplementary Fig. 2b). Similarly, other SINEs could inhibit CRISPR-Cas9 activity across different genomic sites (Supplementary. 2c). Next-generation sequencing (NGS) analysis showed that treatment with SINEs at a therapeutic concentration of 0.5 μM^49^ exhibited modest inhibition of CRISPR-Cas9 activity at *HBB* and *RUNX1* genes in HEK 293T cells (Fig. 1d). Thus, subsequent experiments with SINEs were performed at 0.5 μM concentration.

### SINEs indirectly inhibit Cas9 activity by interfering with the nuclear export of Cas9 mRNA

We next sought to explore the mechanism of action of SINEs for Cas9 inhibition. Surprisingly, it was found that KPT330 at a high concentration of 10 μM showed very limited inhibition of the activity of purified Cas9 protein in an *in vitro* DNA cleavage reaction (Supplementary Fig. 3a). This result suggested that KPT330 as well as other SINEs might inhibit CRISPR-Cas9 activity in an indirect manner. To test this hypothesis, we examined the effects of KPT330 on the cellular activity of Cas9-sgRNA ribonucleoproteins (RNPs). T7E1 analysis showed that 0.5 μM KPT330 exhibited very limited inhibition of Cas9-sgRNA RNP-mediated genome editing at *EMX1* and *AAVS1* sites in Hela cells (Supplementary Fig. 3b). Comparison of the effects of SINEs on transiently transfected plasmids (Fig. 1d) and on RNP of CRISPR-Cas9 (Supplementary Fig. 3a, b) indicated that SINEs were not direct inhibitors of CRISPR-Cas9 but rather functioned at transcriptional or translational level.

It has been well established that KPT330 exerts its anti-cancer activity by inhibiting nuclear transporter protein exportin-1 (XPO1), also known as CRM1^49^. XPO1 is responsible for the nuclear export of various tumor suppressor proteins and can, in association with adaptor proteins, export a wide range of RNA cargos including messenger RNAs (mRNAs), ribosomal RNA (rRNAs), transfer RNA (tRNAs), small nuclear RNAs (snRNA), microRNA (miRNA) and viral mRNAs^50^. To investigate whether SINEs could interfere with the nuclear export of Cas9 mRNA, we analyzed the effects of 0.5 μM KPT330 on the cytoplasmic expression of Cas9 mRNA following transfection of sgRNA- and Cas9-coding plasmids in HEK293T cells. It was found that 0.5 μM KPT330 treatment reduced Cas9 mRNA in the cytoplasm, independently of Cas9-associated sgRNA (Fig. 2a). Moreover, we evaluated the effects of other SINEs on the expression of cytoplasmic Cas9 mRNA and found that all SINEs reduced cytoplasmic Cas9 mRNA in a dose-dependent manner except KPT276 (Fig. 2b). To confirm SINEs affected Cas9 mRNA by interfering with the transport process rather than affecting the stability of cytoplasmic mRNA, we directly transfected cells with *in vitro* transcribed Cas9 mRNA along with sgRNA to bypass the nuclear export process associated with plasmid-encoded Cas9. It was found that 0.5 μM KPT330 did not significantly inhibit the genome modification events mediated by *in vitro* transcribed Cas9 mRNA (Supplementary Fig. 3c), indicating that KPT330 had little impact on Cas9 mRNA stability. These data collectively demonstrated that the inhibitory activity of SINEs was dependent on the nuclear export of Cas9 mRNA.

**Fig. 2:**
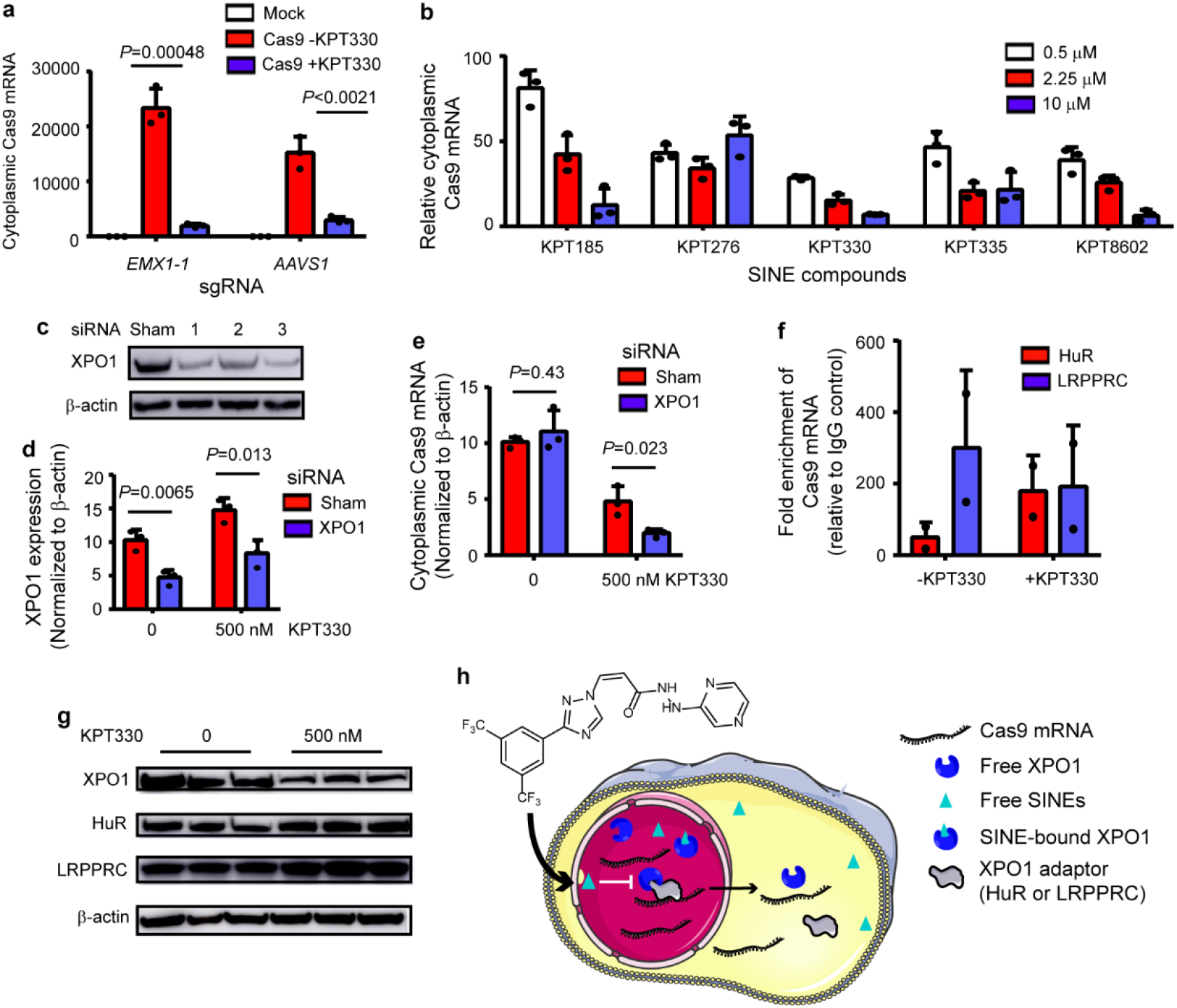
SINEs inhibit the nuclear export of Cas9 mRNA in an XPO1-dependent manner. **a** KPT330 reduces cytoplasmic Cas9 mRNA, as determined by RT-qPCR. Significant difference is determined using Student’s *t* test. **b** SINEs reduce cytoplasmic Cas9 mRNA in a dose-dependent manner, as determined by RT-qPCR. mRNA expression is normalized to drug-free group. For **a-b**, β-actin is used as an internal control. **c** Screening siRNA screen for XPO1 knockdown. **d** Evaluation of the efficiencies of siRNA knockdown of XPO1 expression in the absence and presence of 0.5 μM KPT330. **e** The effects of XPO1 knockdown on Cas9 mRNA transport in the absence and presence of 0.5 μM KPT330. For **d-e**, significant difference between sham and XPO1 knockdown is determined using Student’s *t* test. **f** RIP experiment showing HuR and LRPPRC binding with Cas9 mRNA. **g** The effects of KPT330 treatment on the expression of XPO1, HuR and LRPPRC proteins. **h** Cartoon illustrating SINE-mediated modulation of Cas9 mRNA transport. The above data are shown as mean ± SD (*n* = 2 or 3).

### XPO1 was involved in SINE-mediated inhibition of Cas9 mRNA nuclear export

The cellular target of SINEs, XPO1, is an extensively characterized transporter protein for nuclear protein and RNA cargos. It is therefore possible that SINEs affect the nuclear export of Cas9 mRNA in an XPO1-dependent manner. In the presence of 0.5 μM KPT330, XPO1 knockdown by siRNA (Fig. 2c, d) further reduced cytoplasmic Cas9 mRNA (Fig. 2e), suggesting that XPO1 was involved in SINE-mediated inhibition of Cas9 mRNA transport. It was noted that XPO1 siRNA along did not impact Cas9 mRNA transport (Fig. 2e), likely due to the residual XPO1 (Fig. 2c, d) or alternative transport pathway.

It has been reported that XPO1 exports nuclear mRNA via adaptor proteins human antigen R (HuR) or leucine-rich pentatricopeptide repeat protein (LRPPRC)^50^. RNA immunoprecipitation (RIP) experiments confirmed that HuR or LRPPRC could interact with Cas9 mRNA that was produced from Cas9-encoding plasmid both in the absence or presence of KPT330 (Fig. 2f). Importantly, KPT330 treatment reduced the expression of XPO1 but not HuR or LRPPRC (Fig. 2g), suggesting that SINEs affected Cas9 mRNA transport by targeting XPO1 rather than adaptor protein HuR or LRPPRC. Collectively, these results have demonstrated that SINEs regulated the nuclear export of Cas9 mRNA via XPO1/HuR or XPO1/LRPPRC pathway (Fig. 2h).

### SINEs improve the specificity of Cas9-mediated genome editing in human cells

Next, we sought to establish SINEs as modulatory compounds for controlling the cellular activity of CRISPR-Cas9 tools at the mRNA transport level. We analyzed the effects of 0.5 μM SINEs on the on- and off-target editing of CRISPR-Cas9 at *HBB* and *RUNX1* sites in human cells (Supplementary Fig. 4a). Importantly, it was found that SINEs improve the specificity of CRISPR-Cas9, as defined by the ratio between on-target and off-target activity (Fig. 3a).

**Fig. 3:**
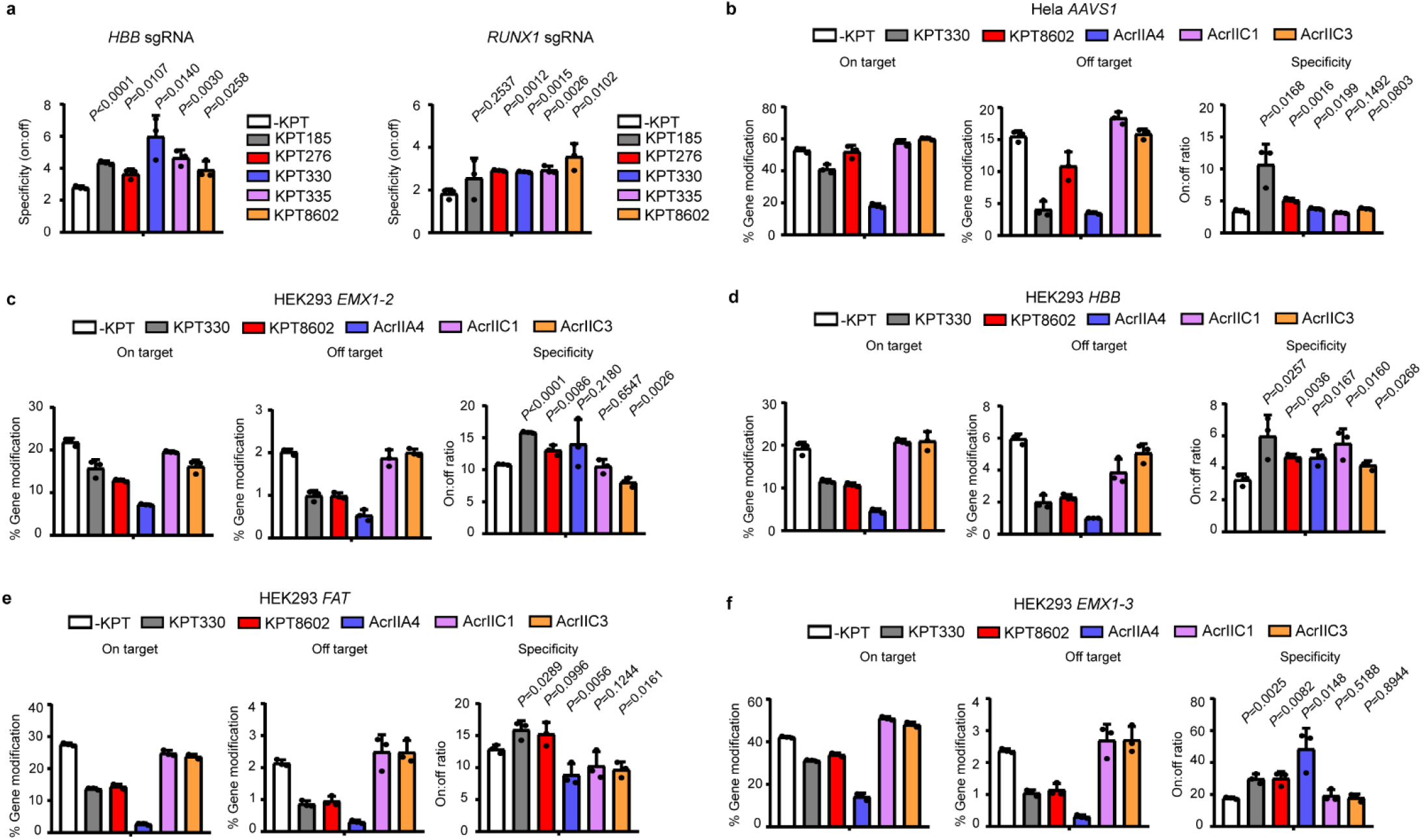
Evaluation of the effects of SINEs on the genome-editing specificity of CRISPR-Cas9. **a** SINEs at 0.5 μM improve the genome-editing specificity of CRISPR-Cas9 at *HBB* gene and *RUNX1* in HEK293 cells. **b** Comparison of the effects of SINEs and Acrs on the genome-editing activity of CRISPR-Cas9 at *AAVS1* site in Hela cells. **c-f** Comparison of the effects of SINEs and Acrs on the genome-editing activity of CRISPR-Cas9 at *EMX1-2*, *HBB*, *FAT* and *EMX1-3* in HEK293 cells. The data are shown as mean ± SD (*n* = 3) and the significant difference between mock and SINE or Acrs treatment is determined using Student’s *t* test.

As previous studies have shown that Acrs can improve the specificity of CRISPR-Cas9^37^, we intended to directly compare SINEs with Acrs for their effects on CRISPR-Cas9 targeting. It appeared that KPT330 and KPT8602 had greatest improvement on CRISPR-Cas9 specificity at *HBB* and *RUNX1* sites respectively. Therefore, we focused subsequent analysis on these two SINE compounds. Meanwhile, strong CRISPR-Cas9 inhibitor AcrIIA4^38^ and weak inhibitors AcrIIC1^32^ or AcrIIC3^32^ were selected as comparison. The comparative experiments were performed across different cell types and genomic sites. We found that KPT330 and KPT8602 treatment could consistently improve the specificity of CRISPR-Cas9 by preferentially inhibiting the off-target activity. Importantly, under all treatment conditions with KPT330 and KPT8602 a considerable fraction of the on-target activity of CRISPR-Cas9 was retained (Fig. 3b-f). By contrast, neither strong or weak Acr inhibitors could consistently improve the specificity of CRISPR-Cas9 (Fig. 3b-f). Although AcrIIA4 appeared to exhibited higher specificity than SINEs at *EMX1-3* site (Fig. 3f), this benefit was associated with markedly compromised on-target activity (Fig 3f and Supplementary Fig. 4b) that prevented the practical applications of AcrIIA4 for modulating CRISPR-Cas9 activity and specificity.

Importantly, KPT330 treatment did not alter the pattern of Cas9-induced mutations, as characterized by the mutation peak upstream of the protospacer adjacent motif (PAM) sequence (Supplementary Fig. 5a). In addition, KPT330 had minor impact on the length (Supplementary Fig. 5b) or frame phase of CRISPR-Cas9-induced indels (Supplementary Fig. 5c). These results suggested that SINEs could modulate the genome-editing activity and specificity of CRISPR-Cas9 without complicating the genomic outcome. It must be noted, however, that the effects of SINEs on CRISPR-Cas9 were dependent on genomic sites, sgRNAs or compounds.

### SINEs improve the editing window of CBE

We next investigated the effects of SINEs on Cas9-derived base editing tools. It has been reported that limiting the exposure of cells to excessive CBEs could improve their editing specificity^13,17^. We hypothesized that the wide targeting window of CBEs is attributed, at least in part, to the uncontrolled activity of base editing agents inside cells, which can be improved by temporally controlling the cellular activity of SINEs. It was found that 0.5 μM KPT330 could inhibit BE3 CBE (rAPOBEC1-nCas9-UGI)-induced C-to-T conversion at various genomic sites (Supplementary Fig. 6a). Similar inhibitory activities of other SINEs were observed with BE3 (Supplementary Fig. 6b) and A3A (hAPOBEC1-nCas9-UGI) (Supplementary Fig. 6c) CBEs. Importantly, KPT185, KPT330, KPT335 and KPT8602 exhibited more inhibition toward out-of-window editing than on-target editing of BE3 or A3A CBEs (Fig. 4a).

**Fig. 4:**
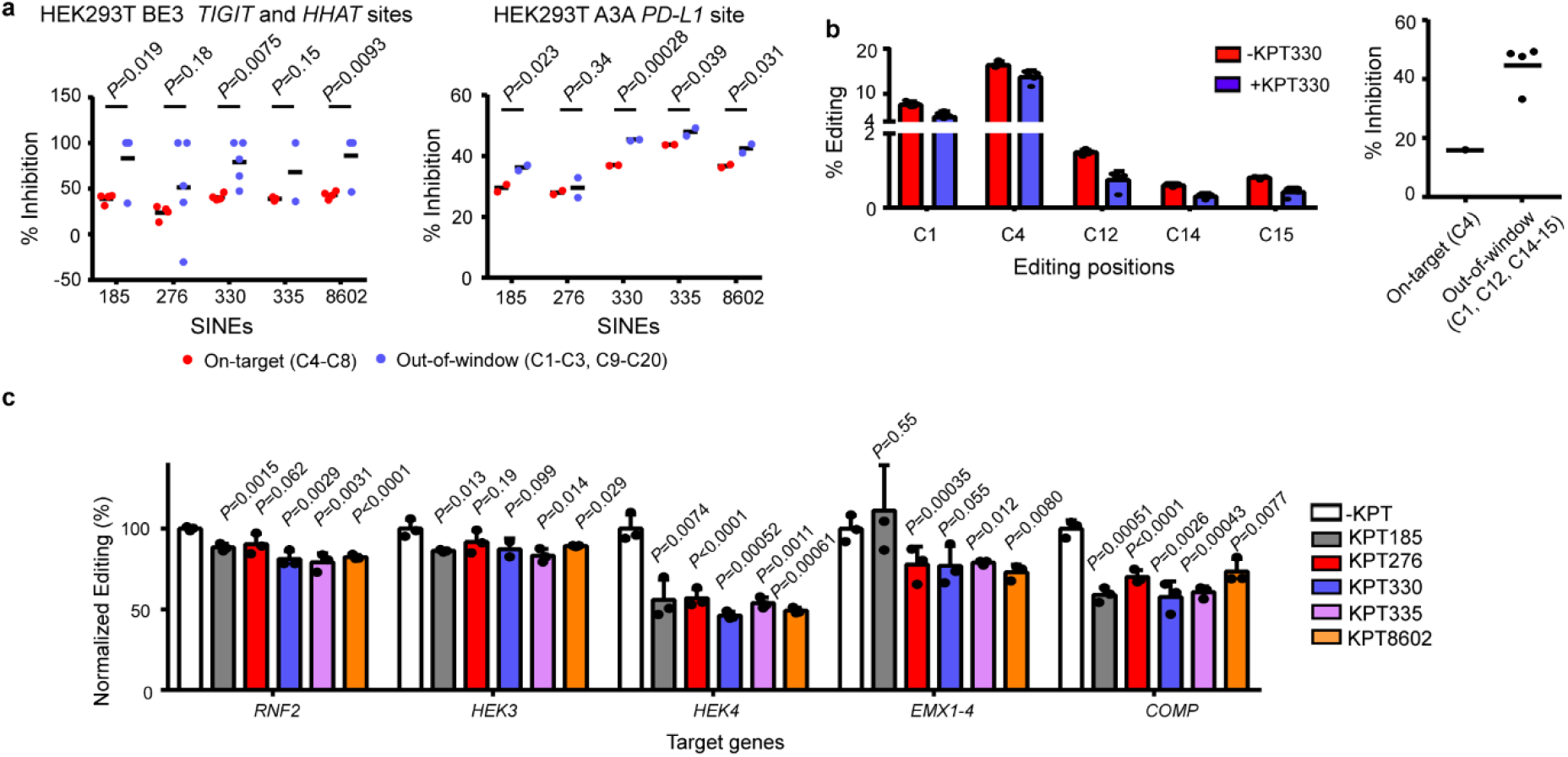
Evaluation of the effects of SINEs on base- and prime-editing tools. **a** The effects of 0.5 μM SINEs on the on-target and out-of-window editing of BE3 and A3A CBEs at different genomic sites. **b** KPT330 at 0.5 μM preferentially inhibits the on-target over out-of-window editing of A3A CBE at the pathogenic *FBN1* site in a HEK293-based disease model of Marfan syndrome. **c** SINEs inhibit the editing efficiency of PE2 and PE3 in HEK293T cells. The data are shown as mean ± SD (*n* = 3) and significant difference between mock and SINE treatment is determined using Student’s *t* test.

The preferential inhibitory activity of SINEs at out-of-window positions allowed a degree of control over the targeting scope of CBEs. To explore the potential application of SINEs during CBE-mediated gene correction, we assessed the effects of KPT330 on A3A CBE editing in a cell-based disease model of Marfan syndrome, where a T7498C mutation was introduced into the *FBN1* gene of HEK293T cells to model the pathogenic C2500R mutation^51^. It was found that KPT330 at 0.5 μM inhibited both on-target and out-of-window editing (Fig. 4b) of A3A CBE but a 2-fold selectivity of inhibition toward out-of-window over on-target editing was observed (Fig. 4b).

The above results suggested that SINEs could inhibit different CRISPR tools carrying the Cas9 module, thus we investigated the effects of SINEs on the recently developed prime editing (PE) tool^52^. Following transfection of PEs, 0.5 μM SINEs were supplemented to cell culture and their effects on PE-induced insertions, deletions and point mutations were analyzed by NGS. It was found that all SINEs showed inhibitory activity toward PEs across different genomic sites in HEK293T cells. The degree of inhibition seemed to be dependent on both the type of SINEs and the edited sites (Fig. 4c). These results again suggested that SINEs functioned as general inhibitors to CRISPR-Cas9, independent of its fusion partners.

## Discussion

Under therapeutic settings, CRISPR-based genome engineering tools are often delivered into human cells as episomal DNA by adeno-associated viruses (AAVs) ^6^. In these applications, CRISPR-Cas9 is associated with off-target effects that are arised from the constitutively active Cas9 protein^7–9^. To overcome these problems, temporal and spatial control strategies have developed to modulate the intracellular activity of CRISPR-Cas9^8,53–56^. For example, inducible promoters have been devised to regulate CRISPR-Cas9 activity at the transcriptional level^57^. Fusion of small-molecule responsive elements to Cas9 proteins allowed the control of CRISPR-Cas9 activity at the post-translational level^54^. In addition, the specificity of CRISPR-Cas9 targeting can be improved by restricting the intracellular half life of Cas9 protein using directly delivered Cas9-sgRNA RNP^18^. These experiments have demonstrated that the intracellular activity of CRISPR-Cas9 can be regulated at multiple levels during the life cycle of CRISPR-Cas9 inside cells.

CRISPR-Cas9 inhibitors provide compelling opportunity to prevent the excessive intracellular activity of CRISPR-Cas9. Despite of the distinct mechanisms of action, most existing CRISPR-Cas9 inhibitors directly intervene the targeting or cleavage processes of CRISPR-Cas9, thus known as direct inhibitors. Specifically, a previous study has established a high-throughput platform to screen for small molecules that can inhibit the *in vitro* cleavage activity of Cas9^45^. Different from this *in vitro* screening system, the present study relied on an EGFP reporter cell-based assay to screen for small molecules that could exert Cas9-inhibiting activity under intracellular conditions. Surprisingly, the best hit of compounds turned out to be an indirect inhibitor that disrupted the nuclear export process of Cas9 mRNA. This compound and its analogues are known as selective inhibitors of nuclear export (SINEs), which are developed to target the nuclear transporter protein XPO1 for anti-cancer treatment. Mechanistic study have revealed that SINE-mediated CRISPR-Cas9 inhibition relies on XPO1 and its adaptor proteins HuR and LRPPRC which are required for mRNA transport. The consistent mechanism of action renders SINEs optimum choice for repurposing for modulating CRISPR-Cas9 activity.

Importantly, one of the identified SINE compounds, KPT330, has been approved to treat relapsed or refractory multiple myeloma (RRMM)^47^ and relapsed or refractory diffuse large B-cell lymphoma (DLBCL)^49^. To the best of our knowledge, KPT330 is thus far the only FDA-approved small molecule that has been identified to bear CRISPR-Cas9-modulating activity. More interestingly, KPT330 as well as other SINEs may improve the genome-editing specificity of CRISPR-Cas9 and the targeting scope of Cas9-derived CBEs. Although these improvements are relatively modest, the comparative analysis has demonstrated that SINEs outperformed traditional Acrs in improving the specificity of CRISPR-Cas9 across different cell types and genomic sites. Furthermore, our discovery provides a new view of manipulating the activity of CRISPR-Cas9 at the mRNA transport level and shed the light on developing next-generation compounds targeting the nuclear export of Cas9 mRNA. In addition, because HuR and LRPPRC are also involved in XPO1-dependent Cas9 mRNA nuclear export, it is possible that the intracellular activity of CRISPR-Cas9 can be regulated by targeting HuR and LRPPRC.

In summary, we report in the present study the discovery of SINEs as the first safe-in-human, indirect inhibitors for CRISPR-based genome editing, base editing and prime editing tools. Our study highlights the importance of Cas9 mRNA transport to the activity and specificity of CRISPR-Cas9. The ability to manipulate Cas9 mRNA transport by SINEs promises a degree of control over the genome editing activity of CRISPR-Cas9. Our discovery should speed advances toward the practical applications of CRISPR-Cas9 technology.

## Online content

Any methods, additional references, Nature Research reporting summaries, Supplementary, supplementary information, acknowledgements, peer review information; details of author contributions and competing interests; and statements of data and code availability are available in the online version of the paper.

## Acknowledgements

J.L. acknowledges the National Natural Science Foundation of China (NSFC, 31600686) and ShanghaiTech University Startup Fund (2019F0301-000-01) for their financial support. J.D. acknowledges NSFC 21672240, NSFC 21421002, Strategic Priority Research Program of the Chinese Academy of Sciences (XDB200203), the Key Research Program of Frontier Sciences from Chinese Academy of Science (QYZDB-SSWSLH-028) and Shanghai Sciences and Technology Committee (18JC1415500 and 18401933502) for their financial support. We thank Prof. Karl Barry Sharpless at Scripps Research and Prof. Kuilin Ding at Shanghai Institute of Organic Chemistry for revision of the manuscript and critical comments. We thank the High-Throughput Screening Platform and at Shanghai Institute for Advanced Immunochemical Studies (SIAIS) at ShanghaiTech University for the support of flow cytometry experiments. We thank Dr. Lichun Jiang and Biomedical Big Data Platform at SIAIS for the support of NGS analyses.

## Author contributions

J.L., J.D. and B.J. conceptualized study. J.L., J.D., S.J.W. and Y.R.C. designed the experiments and analyzed data. T.M, T.G. and G.M. collected and synthesized the compounds in this study. S.-J.W. performed the screen experiments. Y.R.C., S.J.W. and P.Y.H. performed the *in vitro* DNA cleavage assay and CRISPR, BE, PE and RIP experiments in human cells. J.C. analyzed NGS data. J.L. and J.D. wrote the manuscript. All authors discussed the results and commented on and approved the manuscript.

## Additional information

**Supplementary information** is available for this paper online.

**Correspondence and requests for materials** should be addressed to J.L. and J.D.

## Methods

### Cell culture

HEK293T, HEK293-based EGFP reporter and Hela cells were maintained in Dulbecco’s Modified Eagle’s medium (DMEM, ThermoFisher Scientific) supplemented with 10% fetal bovine serum (FBS, ThermoFisher) at 5% CO_2_ and 37 °C in a fully humidified incubator and were passaged when 70-90% confluence was reached. Cell lines were validated by VivaCell Biosciences (Shanghai, China).

### EGFP reporter-based screen for CRISPR inhibitor

HEK293-based EGFP reporter cells (2.5×10^4^ cells/well) were seeded on to 96-well plate. At 24 h after seeding, cells were transfected with 50 ng pST-Cas9 and 50 ng pU6-EGFP T2 plasmids using Opti-MEM medium. At 6 h after transfection, Opti-MEM was replaced with DMEM supplemented with 10% FBS and compounds of indicated concentration. Positive control contains 50 ng pST-Cas9 and 50 ng pU6-EGFP T2 plasmids without compounds. Negative control contains 50 ng pST-Cas9. At 30 h after drug treatment, the fluorescence of cells was quantified by flow cytometry. A cut-off of 80% reduction in the percentage of cells with activated fluorescence was used for the screening experiments.

### Expression and Purification of SpyCas9 proteins

Detailed procedures for SpyCas9 expression and purification can be found in the **Supplementary Information**.

### *In vitro* DNA cleavage and RNP-mediated genome editing

*In vitro* cleavage assay and genome editing using Cas9-sgRNA RNPs were performed as described^46^.

### RT-qPCR quantification of Cas9 mRNA

HEK293 cells (5×10^5^ cells/well) were seeded on to 6 cm plate. At 24 h after seeding, cells were transfected with 2 μg pST-Cas9 plasmids and 1 μg of either pU6-EMX1 T2 or pU6-AAVS1 T2 using Opti-MEM medium. At 6 h after transfection, Opti-MEM was replaced with DMEM supplemented with 10% FBS in the absence or presence of 10 μM KPT330. At 36 h after transfection, nuclear and cytoplasmic RNA were separately extracted (ThermoFisher). RNA of each sample was reverse transcribed into cDNA (Takara). The cDNA was amplified using a SYBR Green Master Mix Kit (Takara) in real-time PCR detection system.

### The effects of SINEs on genome-editing activities of directly delivered Cas9 mRNA

HEK293T cells (2.5×10^5^ cells/well) were seeded on to 24-well plate. At 24 h after seeding, cells were transfected with 3 μg *in vitro* transcribed Cas9 mRNA (APExBIO, R1014) and 2 μg sgRNA by lipofectamine 2000 (Invitrogen). At 6 h after transfection, Opti-MEM was replaced with DMEM supplemented with 10% FBS and 0.5 μM KPT330 is supplemented to cell culture immediately. To analyze genome-editing outcome, cells were harvested at 36 h after transfection and the genomic DNA was extracted, PCR amplified and analysed by T7E1 assay.

### The effects of KPT330 on the genome-editing activity of directly delivered Cas9-sgRNA RNP

Hela cells (2 × 10^5^ cells/well) were harvested, washed with PBS and re-suspended in 20 μL of SE nucleofection buffer (Lonza). Cas9 protein (10 μg) and 15 μg transcribed sgRNA were incubated for 10 min at room temperature to form Cas9-sgRNA RNP complex. Hela cells and Cas9-sgRNA RNP was nucleofected by Lonza 4D nucleofector. Immediately following the nucleofection, 100 μL pre-warmed medium was added into nucleofection cuvettes and the cells were transferred to culture dishes. At 2 h after nucleofection, the cells were attached to the culture dish and 0.5 μM KPT330 was added into cell culture. At 36 h post nucleofection, cells were harvested and genomic DNA was extracted, PCR amplified and analysed by T7E1 assay.

### The effects of SINEs and Acrs on CRISPR-Cas9 activity in human cells

HEK293T, HEK293-based EGFP reporter and Hela cells (2×10^5^ cells/well) were seeded on to 24-well plate. At 24 h after seeding, cells were transfected with 500 ng pST-Cas9 plasmids and 500 ng pcDNA3.1, AcrIIA4-pcDNA3.1, AcrIIC1-pcDNA3.1 or AcrIIC3-pcDNA3.1 plasmid and 250 ng of either pU6-AAVS1 T2, pU6-HBB T2, pU6-VEGFA T2, pU6-IL2RG T2, pU6-RUNX1 T2 or pU6-EMX1 T2, pU6-FAT using Opti-MEM medium. At 6 h after transfection, Opti-MEM was replaced with DMEM supplemented with 10% FBS in the presence or absence of 0.5 μM SINEs of indicated concentrations. Cells were harvested at 36 h after transfection and the genomic DNA was extracted, PCR amplified and analyzed by NGS.

### The effects of SINEs on CRISPR-based BE3 and A3A CBE activities in human cells

HEK293T (2×10^5^ cells/well) were seeded on to 24-well plate. At 24 h after seeding, cells were transfected with 500 ng CBE plasmid and 250 ng sgRNA plasmids using Opti-MEM medium. At 6 h after transfection, Opti-MEM was replaced with DMEM supplemented with 10% FBS in the absence or presence of 0.5 μM SINEs. At 36 h after transfection, genomic DNA was extracted and then analysed by NGS.

### The effects of SINEs on CRISPR-based prime editing in human cells

HEK293T (1×10^5^ cells/well) were seeded on to 48-well plate. At 24 h after seeding, cells were transfected with 750 ng PE2 plasmid, 250 ng pegRNA plasmids and 83 ng nicking sgRNA using Opti-MEM medium. At 6 h after transfection, Opti-MEM was replaced with DMEM supplemented with 10% FBS in the absence or presence of 0.5 μM SINEs. At 72 h after transfection, genomic DNA was extracted, PCR amplified and analysed by NGS.

### RNAi experiments

HEK293T (2×10^5^ cells/well) were seeded on to 12-well plates. At 24 h after seeding, the cells in each well were transfected with 20 pmol siRNA targeting to human XPO1 (GenePharma, Shanghai, China) using lipofectamine 2000 (Invitrogen). At 6 h after transfection, Opti-MEM was replaced with DMEM supplemented with 10% FBS. At 24 h after siRNA transfection, cells were transfected with 1 μg pST-Cas9 plasmids and 0.5 μg pU6-AAVS1 T2 plasmids using lipofectamine 2000 in Opti-MEM medium. At 30 h after siRNA transfection, Opti-MEM was replaced with DMEM supplemented with 10% FBS in the absence or presence of 0.5 μM KPT330. At 30 h after KPT330 treatment, cells were harvested and RNA was fractionated and extracted using RNA extraction kit (ThermoFisher). The cytoplasmic RNA was reverse transcribed into cDNA (Takara) and then amplified using a SYBR Green Master Mix Kit (Takara) in real-time PCR detection system.

### Western blot

Cells were harvested by centrifugation, washed once with PBS and lysed in RIPA buffer (Life Technologies) according to the manufacturer’s instructions. Protein concentrations of cell lysates were determined by bicinchoninic acid (BCA) method according to the manufacturer’ s recommendations (Beyotime Biotechnology). For each sample, 15 μg total protein was loaded and resolved on Bis-Tris 4-12% gels (GenScript) and transferred to a polyvinylidene difluoride (PVDF) membrane using Tris-Glycine Buffer (Sangon Biotech). XPO1, HuR, LRPPRC and β-actin were detected using rabbit anti-XPO1 monoclonal antibody (mAb; Cell Signaling, 46249S, 1: 1,000), anti-HuR mAb (Abcam, ab200342, 1: 1,000) and anti-LRPPRC mAb (Abcam, ab205022, 1: 1,000) and mouse anti-β-actin mAb with HRP conjugate (Cell Signaling, 12262S, 1: 1,000). HRP-linked anti-rabbit IgG antibody (Cell Signaling, 7074P2, 1: 5,000) was used as the secondary antibody. Meilunbio fg super sensitive ECL luminescence reagent (Meilunbio).

### RNA immunoprecipitation (RIP)

HEK293T cells were seeded on to 15 cm dish. At 24 h after seeding, cells were transfected with 40 μg pST1374-Cas9 plasmid at 80% confluence using lipofectamine 2000. At 6 h after transfection, Opti-MEM was replaced with DMEM supplemented with 10% FBS in the absence or presence of 0.5 μM KPT330. At 36 h after transfection, the medium was removed and cells were washed twice with ice-cold PBS and then harvested. RIP experiments were performed Imprint RNA Immunoprecipitation Kit (Sigma-Aldrich, RIP-12RXN). Following manufacturer’s instructions, 3.75 μg LRPPRC (Abcam, ab205022) or 3.75 μg HuR (Abcam, ab200342) antibodies was added into each RIP reaction. IgG (Sigma-Aldrich, M7023) of 3.75 μg was used as a reference. RT-qPCR was used to quantify immunoprecipitation-purified Cas9 mRNA using IgG-enriched mRNA as a reference for non-specific enrichment. The difference of RNA sample preparation was accounted by normalizing the Ct values of antibody-enriched RNA to that of the input RNA in the same RT-qPCR assay. Procedure of calculation is detailed in manufacturer’s instructions. Western blot was performed to rule out the interference of KPT330 treatment on the protein expression of XPO1, HuR and LRPPRC.

### Next-generation sequencing of edited genomic sites

Details for genomic DNA extraction, library preparation, sequencing and analyses can be found in the Supplementary Experimental Procedures.

### Statistical analyses

Two or three biological replicates were performed for each experimental condition. Significant difference was analyzed using two-tailed Student’s *t* test.

### Reporting summary

Further information on research design is available in the Nature Research Reporting Summary linked to this article.

### Data availability

NGS data have been deposited into NCBI SRA database with the accession number PRJNA565327. All other data are available from the corresponding authors on reasonable request.

## References

1 Barrangou, R. et al. CRISPR provides acquired resistance against viruses in prokaryotes. Science 315, 1709–1712 (2007).

2 Jinek, M. et al. A programmable dual-RNA-guided DNA endonuclease in adaptive bacterial immunity. Science 337, 816–821 (2012).

3 Mali, P. et al. RNA-guided human genome engineering via Cas9. Science 339, 823–826 (2013).

4 Cho, S. W., Kim, S., Kim, J. M. & Kim, J. S. Targeted genome engineering in human cells with the Cas9 RNA-guided endonuclease. Nat. Biotechnol. 31, 230–232 (2013).

5 Jiang, F. & Doudna, J. A. CRISPR-Cas9 structures and mechanisms. Annu. Rev. Biophys. 46, 505–529 (2017).

6 Fellmann, C., Gowen, B. G., Lin, P. C., Doudna, J. A. & Corn, J. E. Cornerstones of CRISPR-Cas in drug discovery and therapy. Nat. Rev. Drug Discov. 16, 89–100 (2017).

7 Hsu, P. D. et al. DNA targeting specificity of RNA-guided Cas9 nucleases. Nat. Biotechnol. 31, 827–832 (2013).

8 Pattanayak, V. et al. High-throughput profiling of off-target DNA cleavage reveals RNA-programmed Cas9 nuclease specificity. Nat. Biotechnol. 31, 839–843 (2013).

9 Fu, Y., Sander, J. D., Reyon, D., Cascio, V. M. & Joung, J. K. Improving CRISPR-Cas nuclease specificity using truncated guide RNAs. Nat. Biotechnol. 32, 279–284 (2014).

10 Kosicki, M., Tomberg, K. & Bradley, A. Repair of double-strand breaks induced by CRISPR-Cas9 leads to large deletions and complex rearrangements. Nat. Biotechnol. 36, 765–771 (2018).

11 Li, C. et al. HDAd5/35(++) adenovirus vector expressing anti-CRISPR peptides decreases CRISPR/Cas9 toxicity in human hematopoietic stem cells. Mol. Ther. Methods Clin Dev. 9, 390–401 (2018).

12 Komor, A. C., Kim, Y. B., Packer, M. S., Zuris, J. A. & Liu, D. R. Programmable editing of a target base in genomic DNA without double-stranded DNA cleavage. Nature 533, 420–424 (2016).

13 Rees, H. A. & Liu, D. R. Base editing: precision chemistry on the genome and transcriptome of living cells. Nat. Rev. Genet. 19, 770–788 (2018).

14 Gaudelli, N. M. et al. Programmable base editing of A*T to G*C in genomic DNA without DNA cleavage. Nature 551, 464–471 (2017).

15 Villiger, L. et al. Treatment of a metabolic liver disease by in vivo genome base editing in adult mice. Nat. Med. 24, 1519–1525 (2018).

16 Zeng, J. et al. Therapeutic base editing of human hematopoietic stem cells. Nat. Med. 26, 535–541 (2020).

17 Rees, H. A. et al. Improving the DNA specificity and applicability of base editing through protein engineering and protein delivery. Nat. Commun. 8, 15790 (2017).

18 Kim, S., Kim, D., Cho, S. W., Kim, J. & Kim, J. S. Highly efficient RNA-guided genome editing in human cells via delivery of purified Cas9 ribonucleoproteins. Genome Res. 24, 1012–1019 (2014).

19 Kim, K. et al. Highly efficient RNA-guided base editing in mouse embryos. Nat. Biotechnol. 35, 435–437 (2017).

20 Jin, S. et al. Cytosine, but not adenine, base editors induce genome-wide off-target mutations in rice. Science 364, 292–295 (2019).

21 Zuo, E. et al. Cytosine base editor generates substantial off-target single-nucleotide variants in mouse embryos. Science 364, 289–292 (2019).

22 Bondy-Denomy, J., Pawluk, A., Maxwell, K. L. & Davidson, A. R. Bacteriophage genes that inactivate the CRISPR/Cas bacterial immune system. Nature 493, 429–432 (2013).

23 Pawluk, A., Bondy-Denomy, J., Cheung, V. H., Maxwell, K. L. & Davidson, A. R. A new group of phage anti-CRISPR genes inhibits the type I-E CRISPR-Cas system of Pseudomonas aeruginosa. MBio 5, e00896 (2014).

24 Pawluk, A. et al. Inactivation of CRISPR-Cas systems by anti-CRISPR proteins in diverse bacterial species. Nat. Microbiol. 1, 16085 (2016).

25 He, F. et al. Anti-CRISPR proteins encoded by archaeal lytic viruses inhibit subtype I-D immunity. Nat.Microbiol. 3, 461–469 (2018).

26 Marino, N. D. et al. Discovery of widespread type I and type V CRISPR-Cas inhibitors. Science 362, 240–242 (2018).

27 Pawluk, A. et al. Naturally occurring off-switches for CRISPR-Cas9. Cell 167, 1829–1838 e1829 (2016).

28 Hynes, A. P. et al. An anti-CRISPR from a virulent streptococcal phage inhibits Streptococcus pyogenes Cas9. Nat. Microbiol. 2, 1374–1380 (2017).

29 Uribe, R. V. et al. Discovery and characterization of Cas9 inhibitors disseminated across seven bacterial phyla. Cell Host Microbe 25, 233–241 e235 (2019).

30 Watters, K. E., Fellmann, C., Bai, H. B., Ren, S. M. & Doudna, J. A. Systematic discovery of natural CRISPR-Cas12a inhibitors. Science 362, 236–239 (2018).

31 Cui, Y. R. et al. Allosteric inhibition of CRISPR-Cas9 by bacteriophage-derived peptides. Genome Biol. 21, 51 (2020).

32 Harrington, L. B. et al. A broad-spectrum inhibitor of CRISPR-Cas9. Cell 170, 1224–1233 e1215 (2017).

33 Bondy-Denomy, J. et al. A unified resource for tracking Anti-CRISPR names. Crispr J 1, 304–305 (2018).

34 Lee, J. et al. Potent Cas9 inhibition in bacterial and human cells by AcrIIC4 and AcrIIC5 anti-CRISPR proteins. mBio 9, 02321–18 (2018).

35 Basgall, E. M. et al. Gene drive inhibition by the anti-CRISPR proteins AcrIIA2 and AcrIIA4 in Saccharomyces cerevisiae. Microbiology (Reading) 164, 464–474 (2018).

36 Rauch, B. J. et al. Inhibition of CRISPR-Cas9 with bacteriophage proteins. Cell 168, 150–158 e110 (2017).

37 Shin, J. et al. Disabling Cas9 by an anti-CRISPR DNA mimic. Sci. Adv. 3, e1701620 (2017).

38 Li, J., Xu, Z., Chupalov, A. & Marchisio, M. A. Anti-CRISPR-based biosensors in the yeast S. cerevisiae. J Biol. Eng. 12, 11 (2018).

39 Nakamura, M. et al. Anti-CRISPR-mediated control of gene editing and synthetic circuits in eukaryotic cells. Nat. Commun. 10, 194 (2019).

40 Jiang, F. et al. Temperature-responsive competitive inhibition of CRISPR-Cas9. Mol. Cell 73, 601–610 e605 (2019).

41 Bubeck, F. et al. Engineered anti-CRISPR proteins for optogenetic control of CRISPR-Cas9. Nat. Methods 15, 924–927 (2018).

42 Hoffmann, M. D. et al. Cell-specific CRISPR-Cas9 activation by microRNA-dependent expression of anti-CRISPR proteins. Nucleic Acids Res. 13, e15 (2019).

43 Palmer, D. J., Turner, D. L. & Ng, P. Production of CRISPR/Cas9-mediated self-cleaving helper-dependent adenoviruses. Mol. Ther, Methods Clin Dev. 13, 432–439 (2019).

44 Li, B. et al. Synthetic oligonucleotides inhibit CRISPR-Cpf1-mediated genome editing. Cell Rep. 25, 3262–3272 e3263 (2018).

45 Maji, B. et al. A high-throughput platform to identify small-molecule inhibitors of CRISPR-Cas9. Cell 177, 1067–1079 (2019).

46 Liu, J. et al. Efficient delivery of nuclease proteins for genome editing in human stem cells and primary cells. Nat. Protoc. 10, 1842–1859 (2015).

47 Chari, A. et al. Oral selinexor-dexamethasone for triple-class refractory multiple myeloma. New Engl. J. Med. 381, 727–738 (2019).

48 Kalakonda, N. et al. Selinexor in patients with relapsed or refractory diffuse large B-cell lymphoma (SADAL): a single-arm, multinational, multicentre, open-label, phase 2 trial. Lancet Haematol 7, E509–E522 (2020).

49 Ranganathan, P. et al. Preclinical activity of a novel CRM1 inhibitor in acute myeloid leukemia. Blood 120, 1765–1773 (2012).

50 Natalizio, B. J. & Wente, S. R. Postage for the messenger: designating routes for nuclear mRNA export. Trends Cell Biol. 23, 365–373 (2013).

51 Zeng, Y. et al. Correction of the Marfan syndrome pathogenic FBN1 mutation by base editing in human cells and heterozygous embryos. Mol. Ther. 26, 2631–2637 (2018).

52 Anzalone, A. V. et al. Search-and-replace genome editing without double-strand breaks or donor DNA. Nature 576, 149–157 (2019).

53 Fu, Y. et al. High-frequency off-target mutagenesis induced by CRISPR-Cas nucleases in human cells. Nat. Biotechnol. 31, 822–826 (2013).

54 Davis, K. M., Pattanayak, V., Thompson, D. B., Zuris, J. A. & Liu, D. R. Small molecule-triggered Cas9 protein with improved genome-editing specificity. Nat. Chem. Biol. 11, 316–318 (2015).

55 Frock, R. L. et al. Genome-wide detection of DNA double-stranded breaks induced by engineered nucleases. Nat. Biotechnol. 33, 179–186 (2015).

56 Maji, B. et al. Multidimensional chemical control of CRISPR-Cas9. Nat. Chem. Biol. 13, 9–11 (2017).

57 Gonzalez, F. et al. An iCRISPR platform for rapid, multiplexable, and inducible genome editing in human pluripotent stem cells. Cell Stem Cell 15, 215–226 (2014).

